# Pan-cancer computational histopathology reveals mutations, tumor composition and prognosis

**DOI:** 10.1101/813543

**Authors:** Yu Fu, Alexander W Jung, Ramon Viñas Torne, Santiago Gonzalez, Harald Vöhringer, Artem Shmatko, Lucy Yates, Mercedes Jimenez-Linan, Luiza Moore, Moritz Gerstung

## Abstract

The diagnosis of cancer is typically based on histopathological assessment of tissue sections, and supplemented by genetic and other molecular tests^1–6^. Modern computer vision algorithms have high diagnostic accuracy and potential to augment histopathology workflows^7–9^. Here we use deep transfer learning to quantify histopathological patterns across 17,396 hematoxylin and eosin (H&E) stained histopathology slide images from 28 cancer types and correlate these with matched genomic, transcriptomic and survival data. This approach accurately classifies cancer types and provides spatially resolved tumor and normal distinction. Automatically learned computational histopathological features correlate with a large range of recurrent genetic aberrations pan-cancer. This includes whole genome duplications, which display universal features across cancer types, individual chromosomal aneuploidies, focal amplifications and deletions as well as driver gene mutations. There are wide-spread associations between bulk gene expression levels and histopathology, which reflect tumour composition and enables localising transcriptomically defined tumour infiltrating lymphocytes. Computational histopathology augments prognosis based on histopathological subtyping and grading and highlights prognostically relevant areas such as necrosis or lymphocytic aggregates. These findings demonstrate the large potential of computer vision to characterise the molecular basis of tumour histopathology and lay out a rationale for integrating molecular and histopathological data to augment diagnostic and prognostic workflows.

## Pan-Cancer Computational Histopathology

Computational histopathology algorithms can process and cross-reference very large volumes of data, helping pathologists to navigate and assess slides more quickly and aid in quantifying aberrant cells and tissues^10^. Often based on convolutional neural networks (CNNs), these algorithms build an implicit quantification of histopathological image content, which represent the patterns of the image as seen by the computer. These computational histopathological features are automatically learned for the original task of classifying the entire and/or subregions of images into cancer or non-cancerous tissues. However, once learned, the feature representation may also be used to find similar images^11^ and to quantify associations with traits beyond tissue types^12,13^. This approach, known as transfer learning, has been used to establish associations with genomic alterations^14–19^, transcriptomic changes^20,21^ and survival^22–24^.

Here we performed a pan-cancer computational histopathology (PC-CHiP) analysis to assess the utility of computer vision and transfer learning across 28 cancer types to study the associations of histopathology and genomic driver alterations, whole transcriptomes and survival. At its core, PC-CHiP is based on Inception-V4^25^, an established CNN, which was used to extract a set of 1,536 image features for tissue classification and transfer learning (**Figure 1a**). The algorithm was fine-tuned on 17,396 H&E stained fresh-frozen tissue image slides from The Cancer Genome Atlas (TCGA)^26^, containing specimens from 10,452 individuals from 28 tumor types and 14 normal tissues with matched genomic, transcriptomic and outcome data. Slides were tiled into more than 14 million 256µm x 256µm-sized tiles with a digital resolution of 512 by 512 pixels. Tiles from 9,754 slides with tumor purity greater than 85% were split into 80% training and 20% validation data for training based on 42 broadly defined tissue labels. The resulting quantitative histopathology representation accurately discriminated different tissues, displayed pervasive associations with underlying genomic alterations, wide-spread correlations with transcriptomic signatures and prognostic information across most cancer types.

**Figure 1.**
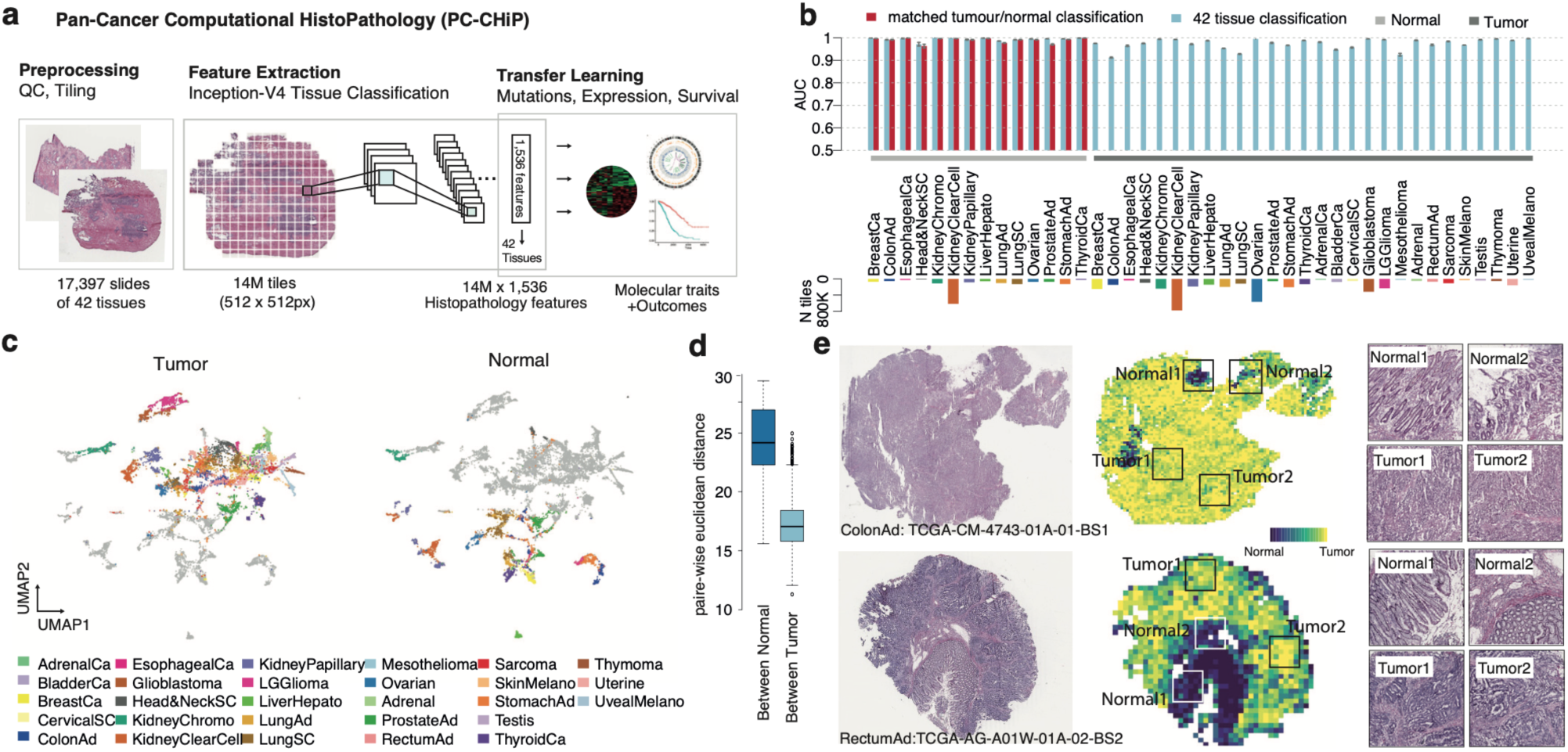
Pan-cancer computational histopathology quantifies tissue-specific morphology. **a**, The PC-CHiP transfer learning workflow for molecular and outcome predictions based on fresh-frozen H&E-stained tissue sections. **b**, Held-back classification accuracy (AUC, upper panel) of 42 tissue types including 18 normal (marked in light gray) and 28 tumour (marked in dark gray). Discrimination of 42 tissues is shown in blue and organ-specific tumor/normal classification in red, with 95% bootstrap confidence intervals each. The lower panel shows the number of tiles in the training dataset for each tissue type. **c**, UMAP dimensionality reduction representation of the 1,536 histopathological features from *n*=100 randomly selected tiles. Tumors are highlighted and coloured on the left and normal tissues on the right. **d**, Pairwise Euclidean distance in the 1,536-feature space for normal tissues (left) and tumor tissues (right). Boxplots depict the quartiles and median, whiskers extend to 1.5× the inter quartile range. **e**, Examples of spatial tumor/normal tissue prediction. For each row, the original H&E-stained slide is shown on the left, the predicted tumor probability of each tile in the middle, and enlarged images of 4 regions indicated in the middle figure are shown on the right side. See **Supplementary Table 2** for cancer type abbreviations.

## Accurate pan-cancer tissue classification and spatial deconvolution

For 14 cancers with normal and tumor images, the average tumor/normal tissue classification AUC was 0.99 (all values given for the held-back validation set; range 0.96 to 0.99, **Figure 1b**). The average pan-tissue AUC of discriminating all 42 different tissues was 0.98 (range 0.91 to 0.99), including the 14 cancer types without matched normal samples (**Supplementary Table 1**; potential limitations are discussed at the end of the manuscript). To achieve this classification, PC-CHiP builds an image representation of each tile consisting of the output of the last 1,536 neurons of the network (**Figure 1a**). Hereafter we will refer to this output as *computational histopathological features* and demonstrate that this representation enables us to derive quantitative associations with a range of molecular traits. As the network was trained to discriminate different tissues, a two-dimensional UMAP representation (**see Methods**) of the computational histopathological features shows clusters corresponding to each tissue class with a certain resemblance of cancers from related organ sites (**Figure 1c, Extended Data Figure 1**). Generally, tumors tend to cluster together, indicating a convergent histological phenotype, usually characterised by a high cell density and loss of tissue architecture – opposed to normal tissues, which are spread over the periphery in the reduced, but also in the original feature space (**Figure 1c,d**).

PC-CHiP was trained using the consensus pathologist estimate of tumour content as soft labels for all tiles of a given slide. While this assigns the same training values for each tile on a given slide, remarkably the network is capable of identifying which tiles on a given slide correspond to cancer and normal regions (**Figure 1e**). This reflects the fact that, based on the comparison of millions of tiles, the algorithm recognises that normal tiles from a tumor slide bear greater similarity with normal slides from the same organ site. Automatic deconvolution achieves a notable accuracy, with an average correlation between algorithm and pathologist-estimated tumor purity equals to 0.26 (range from 0.07 for cervical cancer to 0.6 for uveal melanoma; **Extended Data Figure 2**). The algorithm’s ability to localise signals will be particularly useful when studying the nature of molecular and prognostic associations, as shown in the following sections.

## Histopathological predictions of whole genome duplications

Transfer learning describes the process of using PC-CHiP’s 1,536-dimensional histopathological feature representation to discover novel associations with genomic, transcriptomic and prognostic traits. This amounts to using high-dimensional regression approaches, evaluated by 5-fold cross validation with patient level splits for each fold and separate assessment of each cancer type (**Methods**).

Whole genome duplications (WGD) occur in about 30% of solid tumors, leading to cells with a nearly tetraploid genome, likely as a result of a single failed mitosis^27^. WGD status could be predicted for 19 out of 27 informative cancer types (5-fold cross validated AUC > 0.5, false discovery rate FDR < 0.1, **Methods)** with an average area under the receiver operating characteristic curve (AUC) of 0.73; four cancer types showed an AUC greater than 0.8 (**Figure 2a,b; Supplementary Table 2**). Remarkably, WGD could be predicted for 18 cancer types (FDR<0.1) by models entirely trained on all but the evaluated cancer type, although with slightly reduced accuracy (average AUC=0.68; **Supplementary Table 2, Extended Data Figure 3a**), thus demonstrating that the histopathological characteristics of WGD are largely independent of the specific tissue type.

**Figure 2.**
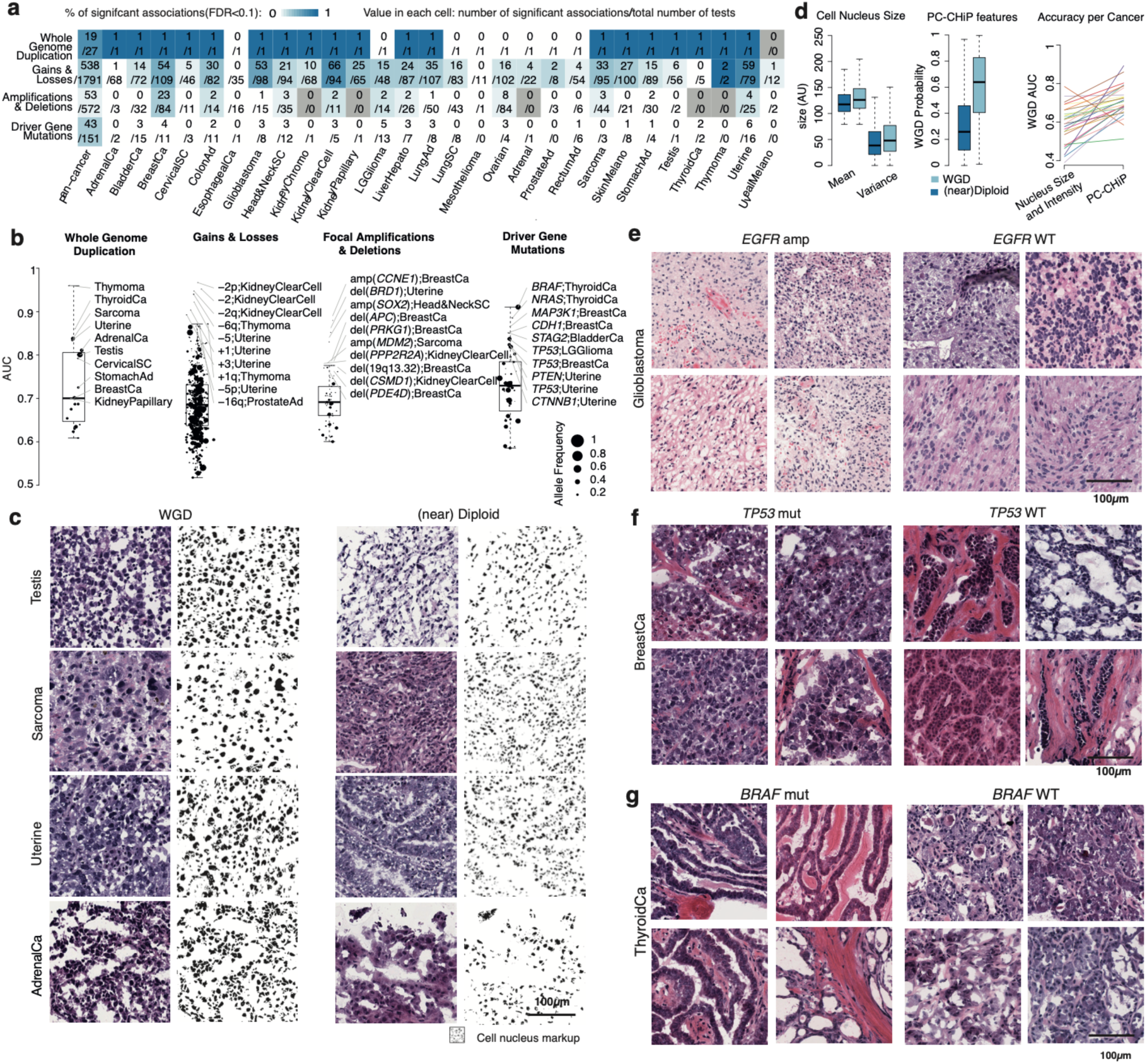
Wide-spread associations between histopathology and genomic alterations. **a**, fraction of significant associations (FDR<0.1) between histopathological features and different types of genetic alterations across cancer types. In each tile the number of significant associations (top) and total (bottom) is given. **b**, Boxplot with AUC for all associations grouped by alteration type indicated on top. Each box corresponds to the distribution of significant associations for each alteration type. **c**, Example tiles with (left) and without (right) whole genome duplication (WGD). From top to bottom, each row corresponds to one cancer type indicated on the left. For each row, haematoxylin and eosin (H&E) stained tumour sections are shown next to a cell nucleus mark-up. **d**, Comparison of traditional hard-coded morphological features nuclear size (right) and CNN-derived predictions between WGD and near diploid samples(middle). The figure on the right side shows the increases of predictive AUC from PC-CHiP compared to hard-coded features. Each line represents one cancer type with the same colors as in **Figure 1. e-g**, Example tiles for indicating presence (left) and absence (right) of particular genomic alterations: *EGFR* amplified glioblastoma (**e**), *TP53* mutant breast cancer (**f**) and *BRAF* mutant thyroid cancer(**g**). All boxplots demarcate the quartiles and median, whiskers extend to 1.5× the inter quartile range.

Tiles with greater probability for WGD displayed an increased nuclear staining, likely due to the higher nuclear DNA content (**Figure 2c**). Explicitly quantifying cell nucleus sizes and intensities using Cell profiler^12^ showed that the average cell nucleus size, as well as its variation across the tumour^28^, was elevated in WGD samples, but these features provided a lower predictive accuracy in all but one cancer type compared to PC-CHiP (average AUC=0.58, range 0.37–0.78; **Figure 2d, Extended Data Figure 3b**). Also, conventional tumour histopathological subtypes and histopathological grade provided a much lower predictive accuracy (average AUC=0.59 for both). This demonstrates how deep learning can automatically detect relevant morphological characteristics, without the need to pre-specify the nature of the patterns to assess, which can lead to more accurate predictions.

## Histopathological associations with copy number alterations

The same approach revealed frequent associations between histopathological patterns and gain and losses of whole chromosomes or chromosome arms (**Figure 2a, Supplementary Table 2**). Yet only subtle histopathological differences between individual aneuploidies were found, and a common feature was high tumour grade, which appears to be a hallmark of chromosomal instability (**Extended Data Figure 4**). The most frequently associated gain was +8q, identified in 12 different cancer types. Conversely, loss of chromosome arm 17p, which harbours the *TP53* tumor suppressor gene, was identified in 12 cancers (**Figure 2e, Extended Data Figure 4b**).

Focal copy number alterations occur on the scale of several megabases and are thought to lead to oncogene amplification and tumor suppressor gene deletions. Based on a catalogue of 140 variants (70 amplifications and deletions each)^29^, 53/563 (9%) alteration:cancer pairs with more than 10 recurrences, showed significant histopathological associations, including 10 amplifications and 43 deletions (FDR < 0.1; **Figure 2a,b; Supplementary Table 2**). The cancer type with the largest number of significant focal copy number alterations was breast invasive carcinoma (4 amplifications and 19 deletions). Notable examples include deletion of *RB1* (13q14.2, AUC=0.75, CI=[0.67, 0.83]) and deletion of *PTEN* (10q23.31, AUC=0.69, CI=[0.63, 0.76]), for which predictions based on histopathological subtypes was inferior (AUC=0.67, CI=[0.62, 0.72] for *RB1* and AUC=0.56, CI=[0.46, 0.66] for *PTEN*).

Recurrently detected focal deletions involve *CSMD1* (8p23.2, 7 cancers) and *PPP2R2A* (8p21.2, 7 cancers; **Supplementary Table 2**). As these 2 deletions frequently co-occur (Cohen’s κ=0.88), it is possible that these histological associations reflect the same underlying alteration. Focal amplification of *EGFR* (located at 7p11.2) in glioblastoma (AUC=0.74, CI=[0.69, 0.78]), was characterised by a distinct small cell morphology of *EGFR-*amplified cancer cells (**Figure 2f**). This histopathological association has been noted previously^30^, although *EGFR* amplifications do not exclusively define a molecular glioblastoma subtype^31^.

## Driver gene mutations

Many oncogenic mutations are point mutations in cancer driver genes. Among all driver genes and cancers tested, 43/151 (28%) gene:cancer displayed significant histopathology associations involving 29 genes (FDR < 0.1, **Figure 2a, Supplementary Table 2**). Interestingly, driver mutations in *TP53*, the most frequently mutated gene in cancers, could be detected in 12/27 (44%) cancer types, including low grade glioma (AUC = 0.84, CI=[0.80, 0.88]), breast invasive carcinoma (AUC = 0.82, CI=[0.78, 0.87]), and uterine cancer (AUC = 0.80, CI=[0.73, 0.87]; **Figure 2b**). Tumours with *TP53* mutations were generally less differentiated and showed higher grade (**Figure 2g, Extended Data Figure 4c**).

A highly accurately predicted cancer driver gene was *BRAF* in thyroid tumors with an AUC as high as 0.92 (CI=[0.87, 0.96]), seemingly associated with a papillary morphology (**Figure 2g**). While it is known that the prevalence of *BRAF* mutations is only about 25% in follicular thyroid carcinomas as opposed to 75% in classical papillary and tall cell thyroid carcinomas^15,32^, these data indicate that the canonical histopathological classification may be further improved (AUC=0.81, CI=[0.69, 0.93]). A similar association was observed for *PTEN* in uterine cancers with an AUC of 0.82 (CI=[0.76, 0.89], **Extended Data Figure 4d**), in part due to the enrichment of *PTEN* mutations in endometrial cancer^33^. When combined with histopathology subtypes, the AUC for *PTEN* mutations in uterine cancer increases to 0.92 (CI=[0.85-1]). These wide-spread associations between genomics and histopathology illustrate how alterations either change cellular morphology or occur preferentially in a histopathologically distinct cellular context.

## Transcriptomic associations reflect tumour composition and proliferation

Associations between gene expression and histopathology may not only reflect distinct tumor cell types with different morphological features, but also stromal and infiltrating immune cells. Overall, we found that 42% of all the gene:cancer pairs tested showed an association between bulk transcriptome and histology across all cancer types (5-fold cross validated *ρ* > 0.25, FDR < 0.1, **Figure 3a, Supplementary Table 2**). For 6% of gene:cancer pairs a correlation *ρ* > 0.5 was found, and 0.2% displayed *ρ* > 0.75. No obvious mechanistic insights were provided by the highest scoring genes, yet many associations correlated with the extent of normal tissue. Interpretable trends emerged at the level of gene sets, showing that genes were enriched in pathways related to the immune system (24 cancers, *n*=106 pathways), followed by cell cycle (*n*=76, 11 cancer types) and signal transduction (*n*=59, 20 cancer types; **Figure 3b**), in broad agreement with recent reports^21^.

**Figure 3.**
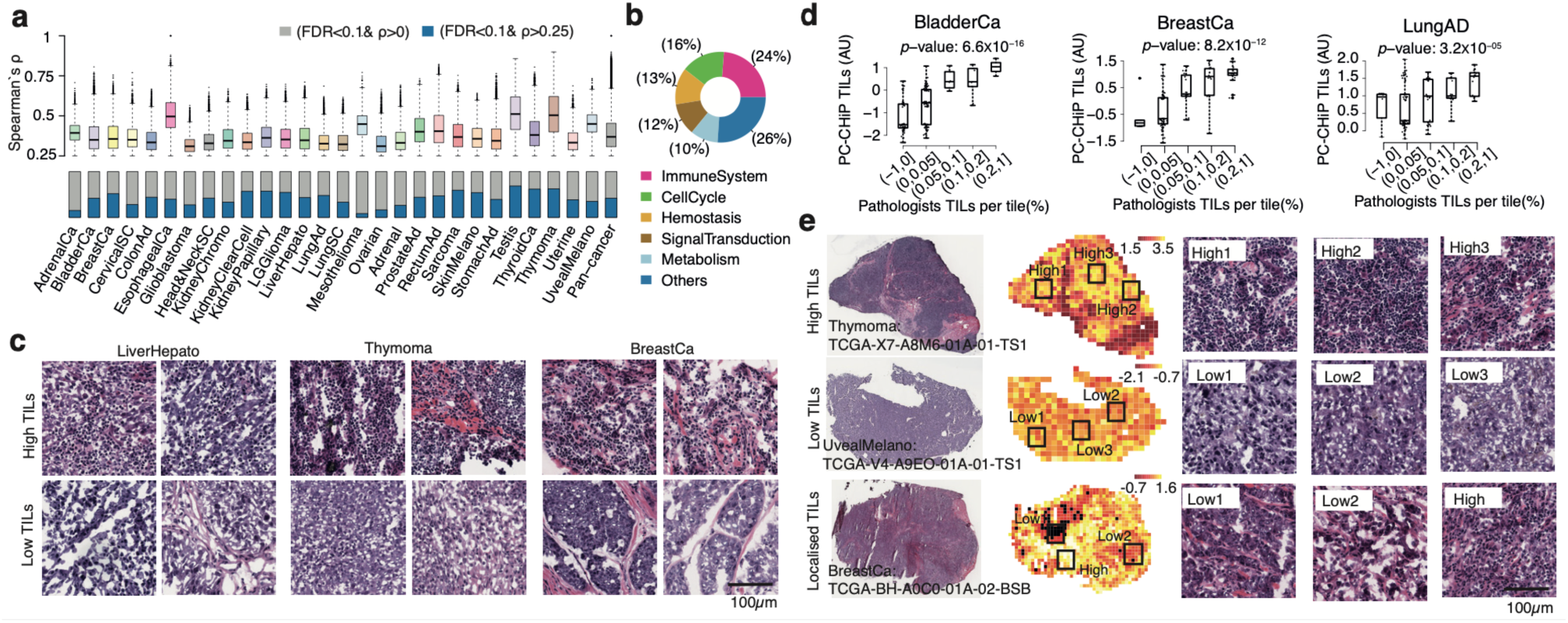
Transcriptomic associations reveal immune infiltration and stromal cell types. **a**, Boxplots of cross-validated Spearman’s rank correlation *ρ* between whole-tumour mRNA expression levels and histopathological features for significant associations (*ρ* > 0.25; FDR < 0.1) across cancer types. The fraction of significant genes is shown in the bottom row. **b**, Gene ontology analysis for significant genes. **c**, Example tiles of predicted TIL-rich (top) and TIL-poor regions (bottom) from hepatocellular carcinoma, breast invasive carcinoma and thymoma. **d**, Systematic blinded assessment of TIL densities by two expert pathologists for three different cancer types. Each box plot shows the predicted TIL scores from PC-CHiP for tiles with different TIL densities, as independently evaluated by pathologists. Boxplots depict the quartiles and median, whiskers extend to 1.5× the inter quartile range. *p*-values for PC-CHiP score from a binomial generalized linear model with TIL fraction as response and added covariate of whole-slide molecular TIL score. **e**, Example slides with high (top row), low (middle row) and localised (bottom row) TILs. From left to right are the original H&E-stained slides, predicted spatial TIL scores and example tiles with high and low TILs. See **Supplementary Table 2** for cancer type abbreviations.

The emerging association between histopathology and cell proliferation was further confirmed by an analysis of transcriptomic proliferation scores^34^, which had histopathologically detectable signals in 25/28 cancer types (*ρ*>0, FDR<0.1, **Supplementary Table 2**). Tumour areas predicted as high proliferation usually had a low stromal content and high grade, while low proliferation overlapped with predicted normal tiles (**Extended Data Figure 5**). This indicates that transcriptomic proliferation scores are in part driven by tumour density and highlights the algorithm’s ability to attribute a molecular association to the presence of histologically normal tissue areas in the tumour section.

## Localisation of immune cells

Tumor infiltrating lymphocyte (TIL) scores^34^ showed a significant *ρ* with histology for all 28 cancer types (*ρ*>0, FDR<0.1), with *ρ*=0.73 (CI=[0.6, 0.82]) for thymoma and notable correlations for breast cancer (*ρ=*0.59, CI=[0.52, 0.64]), bladder cancer (*ρ=*0.58, CI=[0.51, 0.72]) and lung adenocarcinoma (*ρ=*0.48, CI=[0.39, 0.57]; **Supplementary Table 2**). Tiles predictive of TILs indeed contain lymphocytes, which typically are relatively small cells with dark nuclei and scant cytoplasm, often occurring at high densities (**Figure 3c**). These associations were confirmed by a blinded evaluation of tile-level TIL counts and densities by 2 expert pathologists for three representative cancer types (**Figure 3d, Extended Data Figure 6; Methods**). Of note, tile-level predictions were independent of the bulk molecular TIL score confirming the algorithm’s ability to localise the TIL signal to specific areas (*p*=3×10^−3^ for bladder, *p*=6×10^−7^ for breast, and *p*=3×10^−3^ for lung adenocarcinoma; Wald test, including whole-slide TIL score).

The inferred patterns of lymphocytic infiltration, which are recognised prognostic and therapeutic biomarkers^35,36^, were in many cases relatively uniform, with a dispersed distribution of TILs (**Figure 3e**, top and middle panels); in other cases, the signals stemmed from confined regions containing lymphocytic aggregates (**Figure 3e**, bottom panel). Although this approach obviously has many imperfections, it is a remarkable property that these patterns were automatically learned to be a shared feature of tiles from different slides with a high molecular signal of TILs, considering that the algorithm was never provided with defined image sections representing TIL-rich areas.

## Prognostic effects across cancer types

There was a significant association of computational histopathological features with overall survival (OS) in 15/18 cancer types with available data (family-wise error rate FWER<0.05, mean concordance *C*=0.6, range 0.53–0.67; **Methods, Figure 4a, Supplementary Table 3**). Compared to canonical histological subtypes and grades, which are routinely used to assess prognosis, the computational histopathological features showed a significant improvement in 10/16 cancer types. This prognostic signal remained measurable in the majority of these cancer types, even when further including age, gender and tumour stage (**Figure 4b, Extended Data Figure 7, Supplementary Table 3**). As illustrated by the survival curves, PC-CHiP may be used to refine existing stage-based prognosis in breast, head and neck, and stomach cancer as well as, to a lesser extent, clear cell renal cell carcinoma.

**Figure 4.**
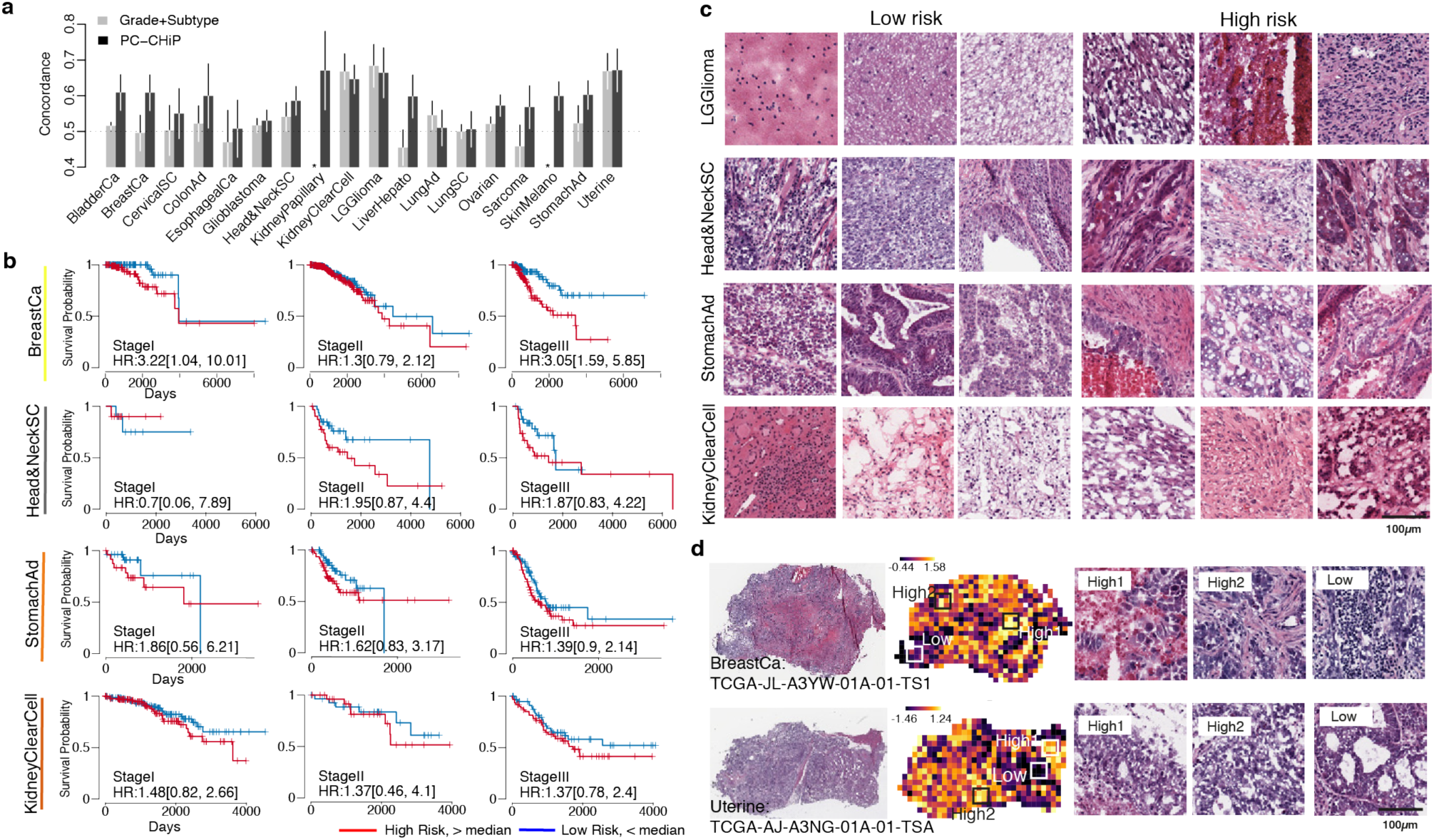
PC-CHiP provides complementary prognostic information. **a**, Predictive accuracy of overall survival using histopathological grade and subtypes (light gray) compared to PC-CHiP (dark gray) in 18 cancers. Each bar corresponds to the mean concordance (with its 95% confidence interval) in 5-fold cross-validation for the dataset and the corresponding cancer type (indicated at the bottom). **b**, Kaplan-Meier plots with high (above median) and low (below median) PC-CHiP risk shown for cancer stage I-III in four different cancer types. **c**, Example H&E-stained tiles with high and low estimated risk based on the histopathological features. Each row corresponds to a cancer type. The first three columns are tiles with low risk and the last three are tiles with high risk. **d**, Risk predictions across whole slides for breast invasive carcinoma and uterine endometrial carcinoma with examples of high and low risk areas enlarged on the right. See **Supplementary Table 2** for cancer type abbreviations.

Reassuringly, many of the prognostic histopathological associations automatically learned by computer vision reflect distinct cancer subtypes as for low grade gliomas (**Figure 4c**). Other features, including necrosis^37^ and high tumour grade, are associated with poor prognosis across tumor types; on the other hand, higher degree of differentiation^38^ and presence of TILs are usually associated with a favourable risk^39^ (**Figure 4c**). Favourable and unfavourable patterns can be frequently identified on the same slide. This highlights the ability of computer vision to deconvolve the content of large tissue sections into molecularly and prognostically distinct areas (**Figure 4d**) with necrosis and lymphocytic aggregates detected on the same specimen. Similarly, areas of low and high grade tumour differentiation identified on the same slide produced favourable and unfavourable risk predictions.

## Validation on external cohorts

The PC-CHiP algorithm exhibited good generalisation on two breast cancer validation cohorts, comprising fresh-frozen H&E stained slides from the METABRIC (*n*=471)^40^ and the BASIS (*n*=151) consortia^41^. The vast majority of genomic associations could be replicated in both cohorts, including associations with *TP53* mutations and WGD with only a moderate drop in accuracy (**Figure 5a,b; Extended Data Figure 8a; Supplementary Table 4**). Similarly, at least 59% (FDR < 0.1) of transcriptomic predictions were recovered in the METABRIC cohort with comparable correlation levels (**Figure 5c; Extended Data Figure 8b**). A subset of both cohorts contained pathologist-evaluated TIL categories, which confirmed the predicted trends, and the algorithm’s ability to localise TILs on a given slide was evident in both cohorts (**Figure 5d, Extended Data Figure 8c**). Prognostic associations were replicated in both cohorts, although at reduced accuracy, particularly in METABRIC (**Figure 5e**). As in the TCGA training data, the algorithm identified necrotic areas and TILs on slides from both cohorts as unfavourable and favourable prognostic markers (**Extended Data Figure 8d**).

**Figure 5.**
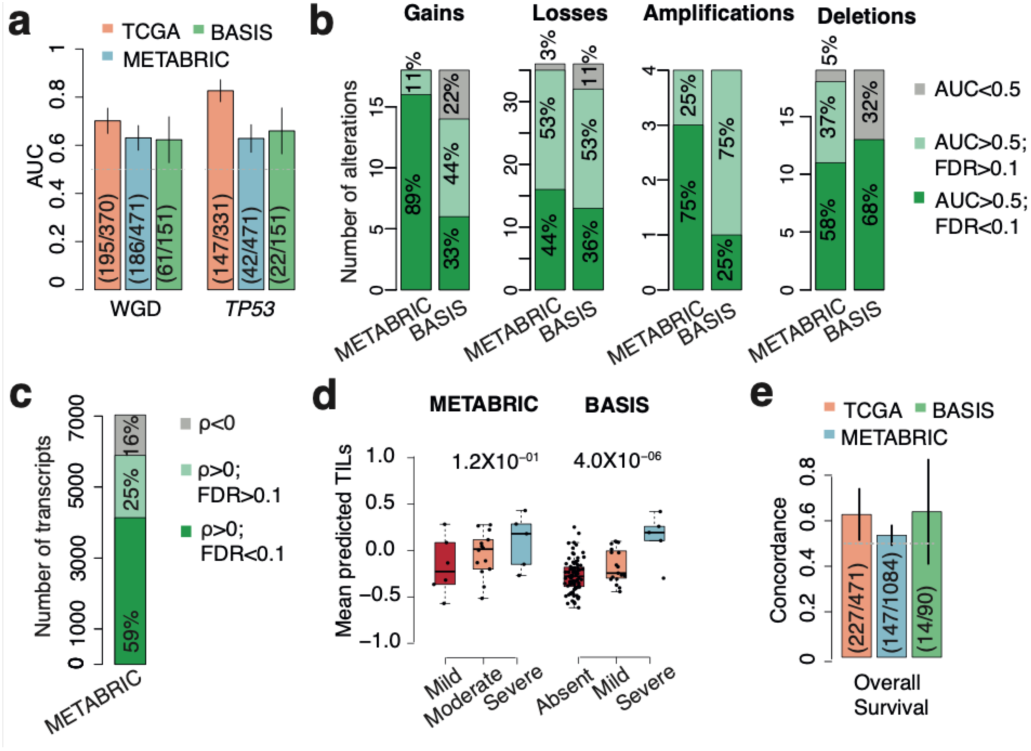
External validation. **a**, Predictive accuracy of WGD, TP53 mutations for the cross-validated predictions in TCGA discovery and METABRIC and BASIS breast cancer validation cohorts. Numbers refer to positive/total cases. **b**, Distribution of validated (deep green), indeterminate (light green) and invalid (gray) associations in METABRIC and BASIS across different alteration types. **c**, Distribution of validated (deep green), indeterminate (light green) and invalid (gray) transcriptomic associations in METABRIC. **d**, Slide-average predicted TIL scores (y-axis) compared to pathology assessment. p-values are for one sided ANOVA. **e**, Prognostic accuracy for overall survival measured by the concordance index C. Numbers refer to events/total cases. All error bars denote 95% confidence intervals.

Of note, there was a considerable tissue misclassification of tiles from the validation cohorts, possibly due to different file formats (**Supplementary Table 4**). Indeed, jpeg quality had a strong confounding effect on histopathological feature representation and tissue classification in the TCGA cohort (**Extended Data Figure 9a-b**). Yet these biases were mostly confined to the initial classification task and were largely mitigated by PC-CHiP’s tissue-aware transfer learning, as confirmed by the high validation rate of molecular associations. Biases of the histopathological feature representation could be further reduced using a file-format aware Inception-V4 architecture and additional data augmentation, which led to a slight drop in the tissue classification accuracy (average AUC=0.95; **Methods, Supplementary Table 4, Extended Data Figure 9c-d**). Transfer learning using histopathological features from the modified architecture produced similar molecular and prognostic associations in TCGA and moderately improved the strength of associations in the METABRIC, but not evidently in the BASIS cohort (**Extended Data Figure 10**).

## Discussion

The results presented here provide a molecular basis for the histopathological observation that tumours are a diverse cellular ecosystem and offer new ways to histologically deconvolute and map its molecularly defined content. Our findings demonstrate that links between a tumour’s morphology and its molecular composition can be found in every cancer type and for virtually every class of genomic and transcriptomic alterations. Whole genome duplications were characterised, and likely caused by nuclear enlargement and increased nuclear intensities, reflecting abnormal chromatin content. Other alterations, such as *EGFR* amplifications or *BRAF* mutations, were associated with a distinct histology and it is currently unknown whether this is a consequence of the alteration or indication that these mutations preferentially occur in particular cell types. Furthermore, a broad range of transcriptomic correlations was found reflecting stromal content, immune cell infiltration and cell proliferation.

In the majority of cancer types, computational histopathological features showed a good level of prognostic relevance, substantially improving prognostic accuracy over conventional grading and histopathological subtyping alone. While it is very remarkable that such predictive signals can be learned in a fully automated fashion, there was no measurable improvement over a full molecular and clinical workup. This might be a consequence of the far-ranging relations between histopathological and molecular phenotypes described here, implying that histopathology is a reflection of the underlying molecular alterations rather than an independent trait. Yet it probably also highlights the challenges of combining histopathological signals from different areas of the same tumour, which requires very large training datasets for each tumour entity.

One of the main current limitations of the study is that the training was performed on fresh-frozen tissue sections. These provide a better preservation of molecular content in comparison to formalin-fixed paraffin embedding, which is the diagnostic standard due to a better preservation of tissue morphology. Also, without further algorithmic amendments, there was a considerable dependence of the histopathological feature representation on image compression algorithms and their parameters, which could in part be mitigated by transfer learning. Given the sensitivity of deep learning algorithms and the associated risk of overfitting, one should generally be cautious about their generalisation properties and critically assess whether a new image is appropriately represented.

While the pervasiveness of associations between histopathology and molecular traits is remarkable, at present they are too weak to replace genetic or transcriptomic tests. However, we expect at least some associations to become more accurate using improved algorithms and larger, ideally spatially annotated, training cohorts. Yet histopathological annotation is possible only for patterns known beforehand and molecular spatial annotation currently has low throughout, but may become possible with spatial transcriptomic^42,43^ and sequencing technologies^44^. Direct training on the molecular trait of interest, or ideally multi-objective learning may provide superior results compared to transfer learning, which risks missing patterns that are irrelevant for the initial classification task. Alternatively, convolutional rather than linear transfer learning has been shown to yield promising results for transcriptomic associations^21^. Also less complex CNN architectures may suffice for classifying histopathology patterns^9,18^, because tumour sections have a defined scale, unlike everyday images.

Looking forward, our analyses reveal the potential of using computer vision alongside molecular profiling. While the eye of a trained human pathologist constitutes the gold standard for recognising clinically relevant histopathological patterns and definitive diagnosis, computers have the capacity to augment these tasks by sifting through millions of images to retrieve similar patterns and establish associations with known and novel traits. Taking the overlays presented in this study as examples, it is not too difficult to imagine computationally augmented and molecularly informed histopathology workflows enabling more precise and faster diagnosis and prognosis in the future.

## Methods

### Images

We collected 17,396 H&E stained histopathology slides of 10,452 patients of 28 broadly defined cancer types from TCGA via the Genomic Data Commons Data Portal^45^ (https://portal.gdc.cancer.gov/), including normal, tumor and metastatic tissue types. Sample inclusion criteria defined by TCGA required primary untreated samples, frozen and sufficiently sized resection samples, and at least 60% tumour nuclei (https://www.cancer.gov/about-nci/organization/ccg/research/structural-genomics/tcga/studied-cancers). Scanned slides usually depict the top and bottom section of the tissue block used for molecular analysis. Only tissue types with at least 50 images with a magnification greater than 20X are included in this study. We first cropped the whole slides into 512 by 512 pixels tiles with 50 pixels overlap at 20X magnification. We then removed blurred and non-informative tiles by filtering on the weighted gradient magnitude (using Sobel operator, tiles with weighted gradient magnitude smaller than 15 for more than half of the pixels were removed). Tiles from tumor samples with tumor purity greater or equals to 85% were used in training/validation to avoid miss labelled tiles in the training process. To avoid bias that is caused by image preparation in individual laboratories, we randomly selected 80% images from each centre for training. In total, we used 6,564,045 tiles from 8,067 slides for training, 1,357,892 tiles from 1,687 slides for validation and 6,641,462 tiles from 7,672 slides for testing.

### Pan-Cancer Computational Histopathology (PC-CHiP)

A pretrained Inception-V4^25^, a deep convolutional neural network, was used to classify tiles into 42 classes and to extract histopathological features from each tile. We applied sample specific label smoothing, an adapted version of the label smoothing method first introduced in Inception-V3^46^ for model regularization, to avoid overfitting. In short, for a sample of tissue *i*, we set ground-truth distribution *q*(*k*) to *q*(*k*) *= p*_*T*_ for *k=i* and *q*(*k*) *=* (*1 – p*_*T*_) */* (*N – 1*) for all ***k ≠ i***, where *N=*42 is the total number of classes and *p*_*T*_ is the tumor purity of the sample. The model was trained in Tensorflow using Slim^47^ with the default hyperparameters for 100K steps (∼1 epoch). The scripts used for training and the retrained model checkpoint can be found on our Github repository.

We retrieved the probability for all 42 classes for each tile and the associated 1,536 histopathological features from the last hidden layer of the trained Inception-V4. As in practice the cancer origin is usually known, we also computed tumor/normal classification within cancer types for each tile by comparing only the probability of being normal or tumor of that cancer type.

To visualise the tiles represented by the 1,536 histopathological features, we applied Uniform Manifold Approximation and Projection for Dimension Reduction (UMAP^48^) for dimension reduction for a subset of tiles (50 tiles were randomly selected from high tumor purity images). As the distances between data points in the original dimension are not preserved in the low dimension generated by UMAP, we also calculated the mean pairwise Euclidean distance between tissue types in their original dimension.

### Algorithmic modifications

In order to reduce the effect of confounding image quality, we modified the classification layers (both the last and auxiliary) for the probability to read *P* = logit^−1^ (*X β* + *Z α*), where *X* ∈ ℝ ^*n* × 1536^ denotes the matrix of PC-CHiP features, *β* ∈ ℝ ^1536 × 42^ being the weight matrix mapping features to the 42 labels, to include the additional term *Z α*, where *Z* ∈ {0,1}^*n* × *f*^ is an indicator matrix of the f confounding factors and *α* ∈ ℝ ^*f* × 42^ being the corresponding weight matrix to absorb the undesired effects of different image qualities and prevent them from being implicitly learned on the deeper layer of the model. Results shown in **Figures 1-4** are based on the generic Inception-V4 architecture with default training parameters and **Figure 5** is based on the modified algorithm, with additional data augmentation to supersede any pre-existing jpeg patterns and heavy color augmentation to overcome systematic differences in H&E staining (random hue rotations by –90 to 90 degrees)^49^.

### Transfer Learning

Regularised generalised linear models, which are broadly analogous to Inception’s original multinomial classification layer, were used to learn molecular associations for each image tile. These models used the set of 1,536 histopathological features and the tissue type encoded as additive indicator variables as predictors and were fitted using the “glmnet” R package^50^. Per slide predictions were calculated by averaging the prediction of all tiles within that slide. To avoid normal contamination, only samples with tumor purity greater than or equal to 85% were included during training. The model performance was reported by the mean predicted accuracy of 5-fold cross validation, split at the level of patients. 100 tiles were randomly selected from each whole slide. Within each fold, 10-fold cross-validation was used to select the glmnet regularisation parameters (folds splitted at patient level). For each of the five folds, a *p*-value was calculated by evaluating model predictions on the held-back fifth using Wilcox’s rank sum test for categorical predictions of genomic data (equivalent to using AUC as a readout), or Spearman’s rank correlation test^51^ for quantitative predictions of transcriptomic data. The resulting 5 *p*-values from each test were combined into a single *p*-value statistic using Fisher’s method to assess whether there was a measurable level of association across folds^52^. Combined *p*-values were then adjusted to control the False Discovery Rate (FDR) across the entirety of cancer:alteration pairs tested using the method of Benjamini and Hochberg^53^. Confidence intervals for the average AUC across folds were estimated using the “cvAUC” R package^54^. Average rank correlations *ρ* and the corresponding confidence intervals were calculated using a tanh^-1^ Fisher transformation^55^. Predictive accuracy was evaluated within each cancer type to avoid reporting associations driven by different prevalence and levels of molecular traits across cancer types. Example tiles and slide overlays shown in this study were from held back validation folds.

### Genomic alterations

Point mutations (single nucleotide variants and short deletions and insertions) were called using CaVEMan and pindel algorithms plus a set of dedicated post-processing filters as described previously^56^ for 8,769 TCGA patients. Absolute copy number was called using the ASCAT algorithm^57^. Whole genome doubling (WGD) status was determined using the criteria described previously^27^. Chromosome and chromosome arm level gains and losses were retrieved from Ref. ^58^. Focal amplifications and deletions were based on regions defined in Ref. ^29^. For each of the amplified regions, samples with an absolute copy number of at least 10 were called amplified; for each of the deleted regions, only samples with ≤ 1 copy in the absence of WGD and samples with ≤ 2 copies in the presence of WGD were called deleted. We performed LASSO regularised multinomial regression models to classify gain, non-altered and loss of 56 chromosome or chromosome arm. We applied logistic regression with LASSO penalization for dichotomous genomic alterations. Per alteration AUCs were then calculated in a one vs the rest fashion (ex. gain vs. not altered and loss) for each cancer type using the statistical procedures described in the previous section.

### Gene expression

Log transformed upper quantile normalized gene expression from RNA sequencing data was used as a readout. We performed linear regression with LASSO penalization on 17,529 genes that were expressed in at least 60% of the samples. Spearman’s rank correlation *ρ* and the predicted explained variance *R*^2^ was calculated for each gene:cancer pair to evaluate the model performance. Associated *p*-values for *ρ*>0 were estimated by Spearman’s rank correlation test^51^. The *p*-values were then corrected controlling the family-wise error rate (FWER) using the method of Bonferroni. In order to identify functional classes of genes that can be predicted by histopathology features, we then performed gene set enrichment analysis (GSEA)^59^ for a collection of REACTOME pathways^60^. A normalised enrichment score and *p*-value were calculated for each pathway in each cancer type. The *p*-values were corrected to control the FDR. Finally, we performed regression on gene expression based proliferation score and tumor infiltrating lymphocytes signature^34^ using the same method used for single gene expression.

### Prognostic associations

Survival analysis was performed using penalized Cox’s proportional hazard regression^61^ using a mixture of *L*1 and *L*2 regularization, often referred to as Cox elastic net^62^. To evaluate discriminative performance we used Harrell’s *C*-index as a measure of the concordance between predicted and actual risk^63^. In order to obtain a scalable and sparse solution, we deployed proximal gradient descent for our parameter updates^64^. Due to the large-scale nature of the problem, an exhaustive hyperparameter search was infeasible. Therefore, hyperparameters, particular *L*1/*L*2 penalization strength, have been automatically determined using Bayesian optimization^65^. Twenty repetitions of 5-fold cross-validation were used to evaluate model performance. Each fold was further split into a training set (85%) and a validation set (15%).

A total of 6 models, each combining different combinations of variables, were evaluated for specific cancer types from TCGA. Cancers were included in the analysis if the sample size was at least 160 individuals and censorship was less than 90%. The first model (“histology”) contains the histological subtype and the corresponding tumour grade information. Routine clinical information for each individual including age at cancer diagnosis, gender, cancer stage and the histopathology features form the second model (“clinical”). The third model (“clinical + expression”) is a combination of clinical and gene expression data. Model four (“PC-CHiP”) uses the extracted histopathological features from the CNN. Model five (“clinical + PC-CHiP”) contains the histopathology features and the clinical data. Lastly, Model six (“all”) is a set of all covariates. If observations have been missing, particularly for the gene expression data, mean imputation has been applied. The gene expression data comprises the first 30 components of a principal component analysis (PCA). For the survival analysis with the histopathology features, each extracted tile has been used as an individual observation. A global risk estimate is obtained using the average risk across the tiles from a patient.

Three different strategies were employed to assess the value of adding PC-CHiP to models based on conventional variables. First, it was tested whether the cross-validated linear predictor obtained using PC-CHiP alone added significant signal in a multivariate model. Second, it was assessed whether the pretrained predictor based on PC-CHiP improved the concordance *C* in a cross-validation setting. Third, a likelihood boosting approach was used for training Cox models from scratch combining clinical/gene expression data with the histopathology features. To compare predictive performance across models we examined the distribution of concordance indices across folds as well as the mean difference concordance within folds. Furthermore, we used a paired-Wilcoxon sign rank test to compare *C* estimates across models. To account for multiple comparisons we used the Holm-Bonferroni correction as FWER procedure. Survival curves have been estimated using the Kaplan-Meier estimator.

### External validation using the METABRIC and BASIS dataset

H&E stained slides from frozen tissue samples were downloaded and tiled into 512 by 512 pixel tiles at 20X magnification in the same fashion as in TCGA. WGD status was calculated using the methods described previously^66^. The amended Inception-V4 architecture, preprocessing scripts and the retrained model checkpoint can be found on our Github repository. Per slide predictions for METABRIC and BASIS were calculated using all tiles.

### Expert blinded assessment of TIL counts

To assess whether predicted TIL levels reflect the true level of immune cells for individual tiles, we randomly selected 150 tiles with different levels of TILs from 3 cancer types including breast, bladder and lung cancer (each 50 tiles from the highest 10% quantile, 50%-70% quantile and lowest 10% quantile). The number of TILs for each tile were independently evaluated by 2 expert pathologists, which were blinded to the predicted scores. The total number of nuclei was automatically learned using the algorithm described^67^. The relationship of predicted TIL sores from PC-CHiP, slide level transcriptomic TIL scores and the true TILs score was modeled using multiple linear regression.

## Data availability

TCGA data (images, genomic, transcriptomic and clinical data) is freely available from http://gdc.cancer.gov. For METABRIC, images, genomic and transcriptomic data is available at EGA https://ega-archive.org/dacs/EGAC00001000484 through controlled access^40^; clinical data is available from https://www.cbioportal.org/. For BASIS, genomic data is freely available from ftp://ftp.sanger.ac.uk/pub/cancer/Nik-ZainalEtAl-560BreastGenomes; clinical data is published^41^; histopathology images are available as controlled access data at the EGA (in process; link will be inserted at publication).

## Code availability

The computational histopathology algorithm and analysis code are available at https://github.com/gerstung-lab/PC-CHiP.

## Acknowledgements

AWJ and MG are supported by grant NNF17OC0027594 from the Novo Nordisk Foundation. LM is a recipient of a CRUK Clinical PhD fellowship (C20/A20917). The results shown here are in part based upon data generated by the TCGA Research Network: https://www.cancer.gov/tcga. We thank Carlos Caldas, Suet-Feung Chin, Yinyin Yuan and the METABRIC consortium, as well as Mike Stratton, Marc Van de Vijver and the BASIS consortium for assistance and sharing data. We also thank all members of the Gerstung Lab, Iñigo Martincorena and Andrew Lawson for critical comments on the manuscript.

## Author Contributions

YF retrieved and quality controlled all images, developed and trained the deep learning algorithms, performed statistical tests for genomic and molecular association and created all figures. AWJ performed survival analysis, reviewed statistical procedures and applied multiple testing adjustments. RVT and MG extended the Inception-V4 algorithm. SG provided copy number and annotated mutation data. HV extracted mutational signature data. AS performed nuclei segmentation. LY curated validation data. LM and MJL reviewed histopathology images and performed blinded assessment of TILs. MG conceived and supervised the study. YF, AWJ and MG wrote the manuscript with input from LM and all other authors who approved the manuscript.

## Extended Data Figures

**Extended Data Figure 1.**
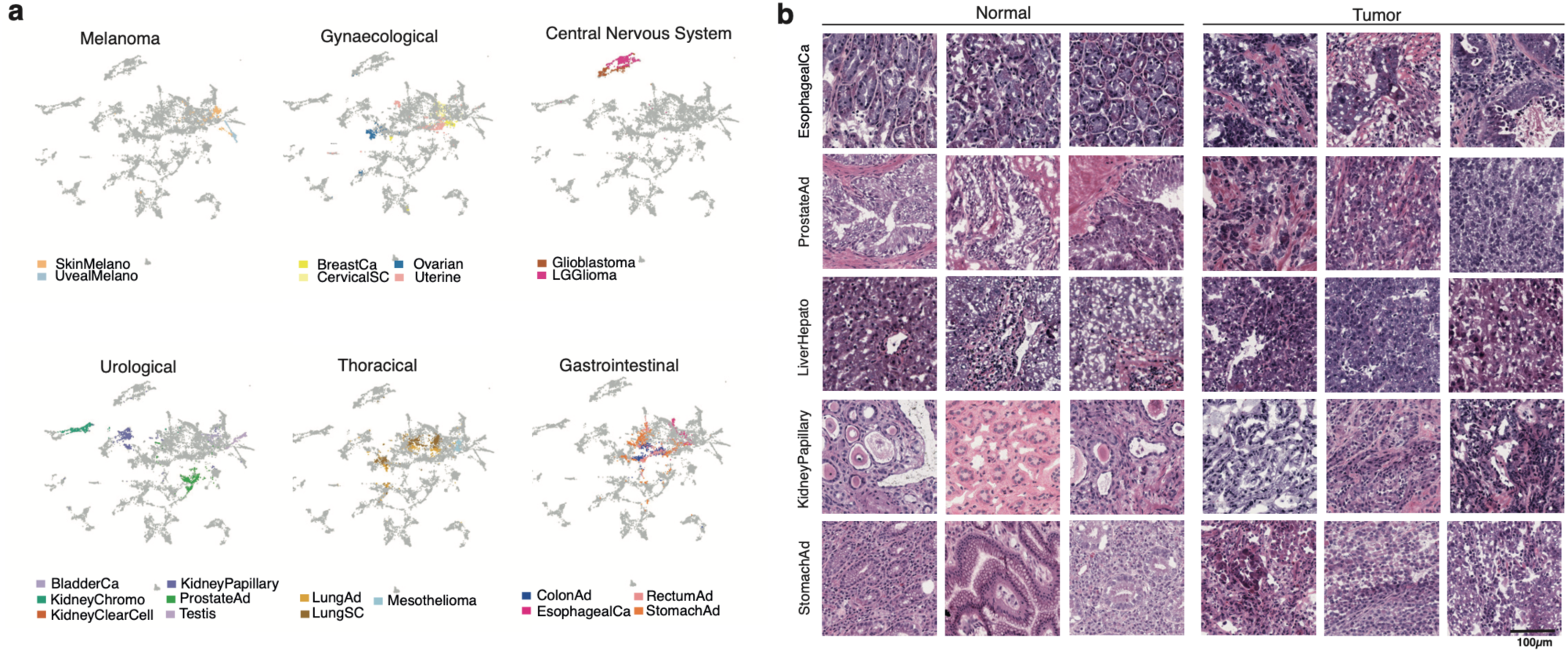
Computational histopathological features discriminate between different tissue types. **a**, UMAP dimensionality reduction representation of the 1,536 histopathological features from randomly selected tiles colored by groups of cancer types. **b**, Example tiles from H&E-stained tissue sections of normal and tumor samples from different cancer types (arranged by row).

**Extended Data Figure 2.**
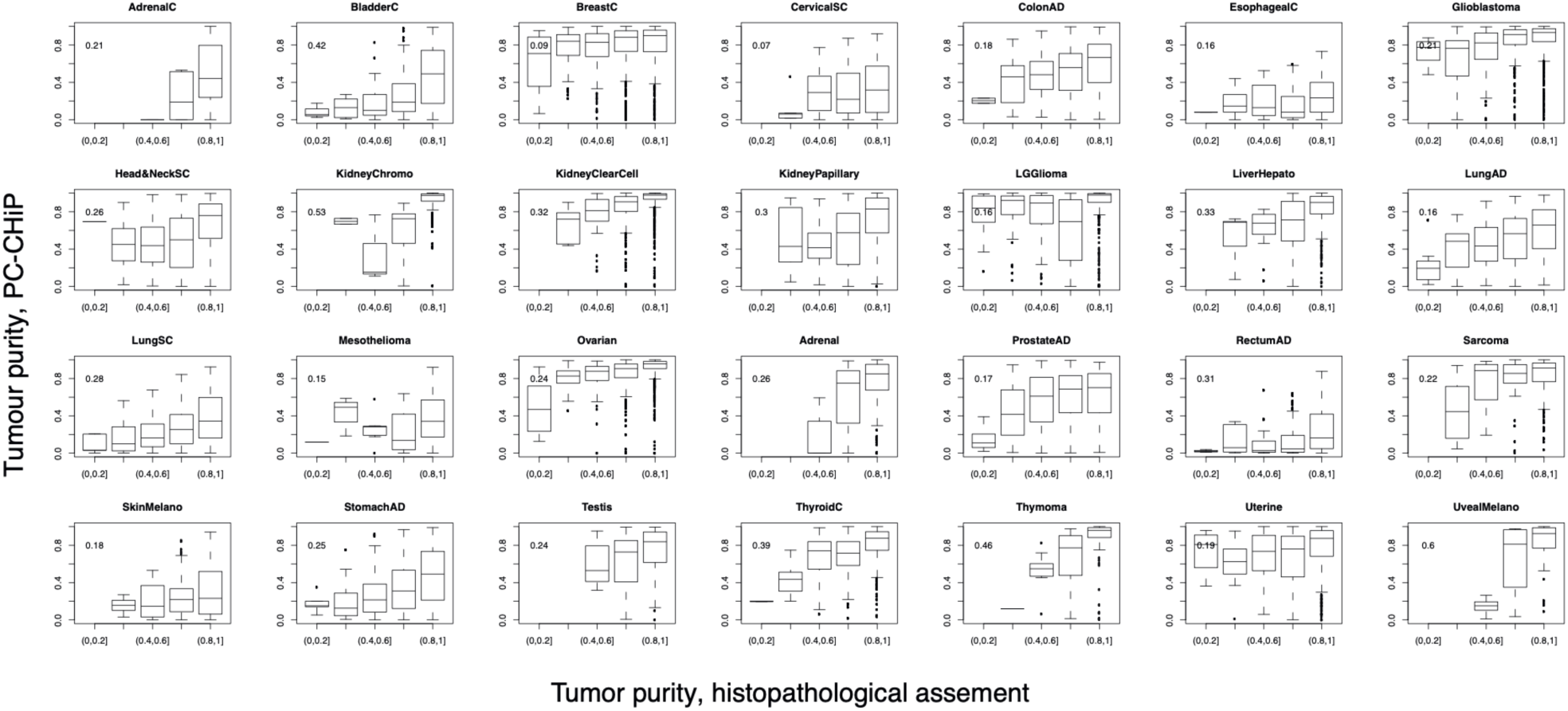
The distribution of predicted tumor purity by histopathological features for samples with different histopathologists evaluated tumor purity. Each boxplot corresponds to one cancer type, each box corresponds to the predicted tumor purity from histopathological features for samples with the histopathologist evaluated tumor purity indicated on x-axis.

**Extended Data Figure 3.**
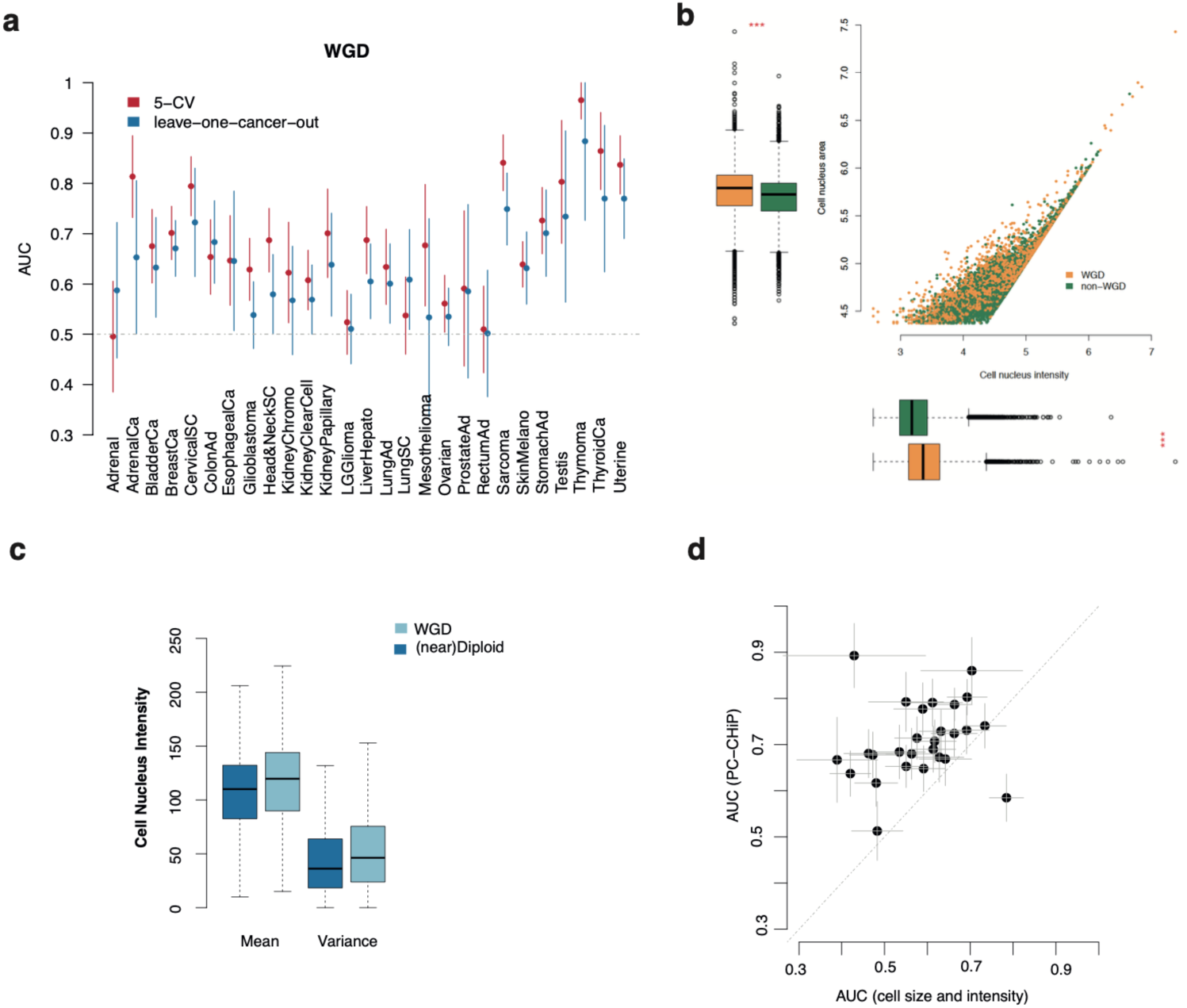
Pan-cancer morphological features of whole genome duplications. **a**, Histopathological prediction of WGD using 5-fold cross validation (red) and models trained leaving out one cancer type (blue). **b**, Distribution of cell nucleus size and intensity of samples with and without WGD. Each dot in the scatter plot corresponds to one of 12,000 tiles that were randomly selected across cancer types. The cell nucleus size and intensity were calculated using Cell Profiler with a pipeline provided by the software provider. Boxplots depict the quartiles and median, whiskers extend to 1.5× the inter quartile range. **c**, barplots show the distribution of the mean and variance of cell nucleus intensity between tiles with and without WGD. **d**, Increases of predictive AUC from PC-CHiP (y-axis) compared to hard coded features (x-axis) for a set of 500 randomly selected tiles for each cancer type. Each dot represents a cancer type. 95% confidence interval estimated from bootstrap.

**Extended Data Figure 4.**
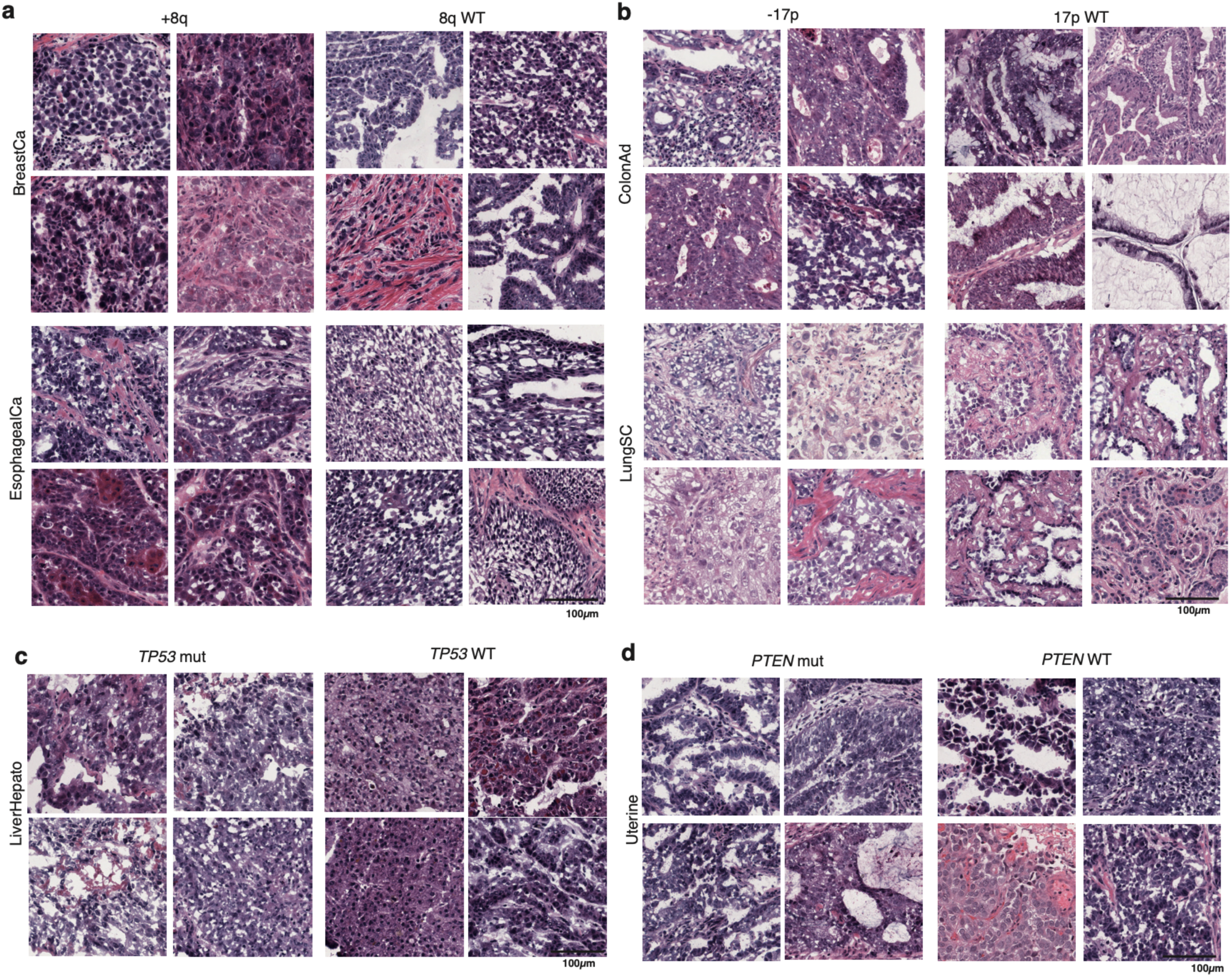
Example tiles for associations between computational histopathological and genomic alterations. **a**, Four randomly selected example tiles for chromosome 8q gain (left column) and wild type (right column) breast invasive carcinoma (top row) and esophageal carcinoma (bottom row). **b**, Example tiles for chromosome 17p loss (left column) and wild type (right column) for colon adenocarcinoma (top row) and lung squamous cell carcinoma (bottom row). **c**, Example tiles for *TP53* mutated (left column) and wild type (right column) liver cancer (hepatocellular carcinomas). **d**, Example tiles for *PTEN* mutation (left column) and wild type (right column) for uterine cancer.

**Extended Data Figure 5.**
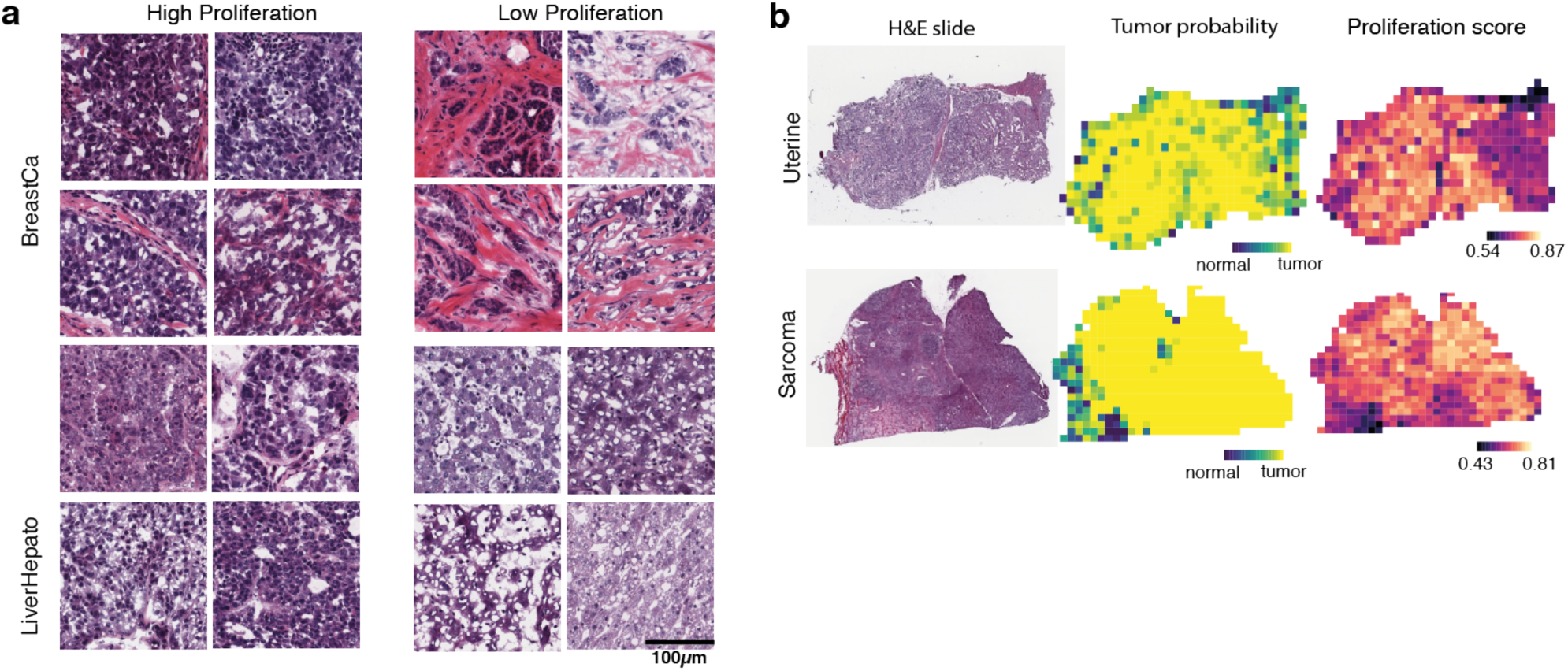
Histopathological associations with transcriptomic cell proliferation scores. **a**, Example tiles for high (left column) and low proliferation (right column) for breast invasive carcinoma (top row) and hepatocellular carcinoma (bottom row). Four randomly selected tiles are shown for each tumour type. **b**, Spatial proliferation predictions for tumor/normal areas within one slide from a uterine cancer (first row) and a sarcoma (second row). From left to right are H&E stained tissue slide, spatial tumor/normal tissue prediction and spatial proliferation predictions.

**Extended Data Figure 6.**
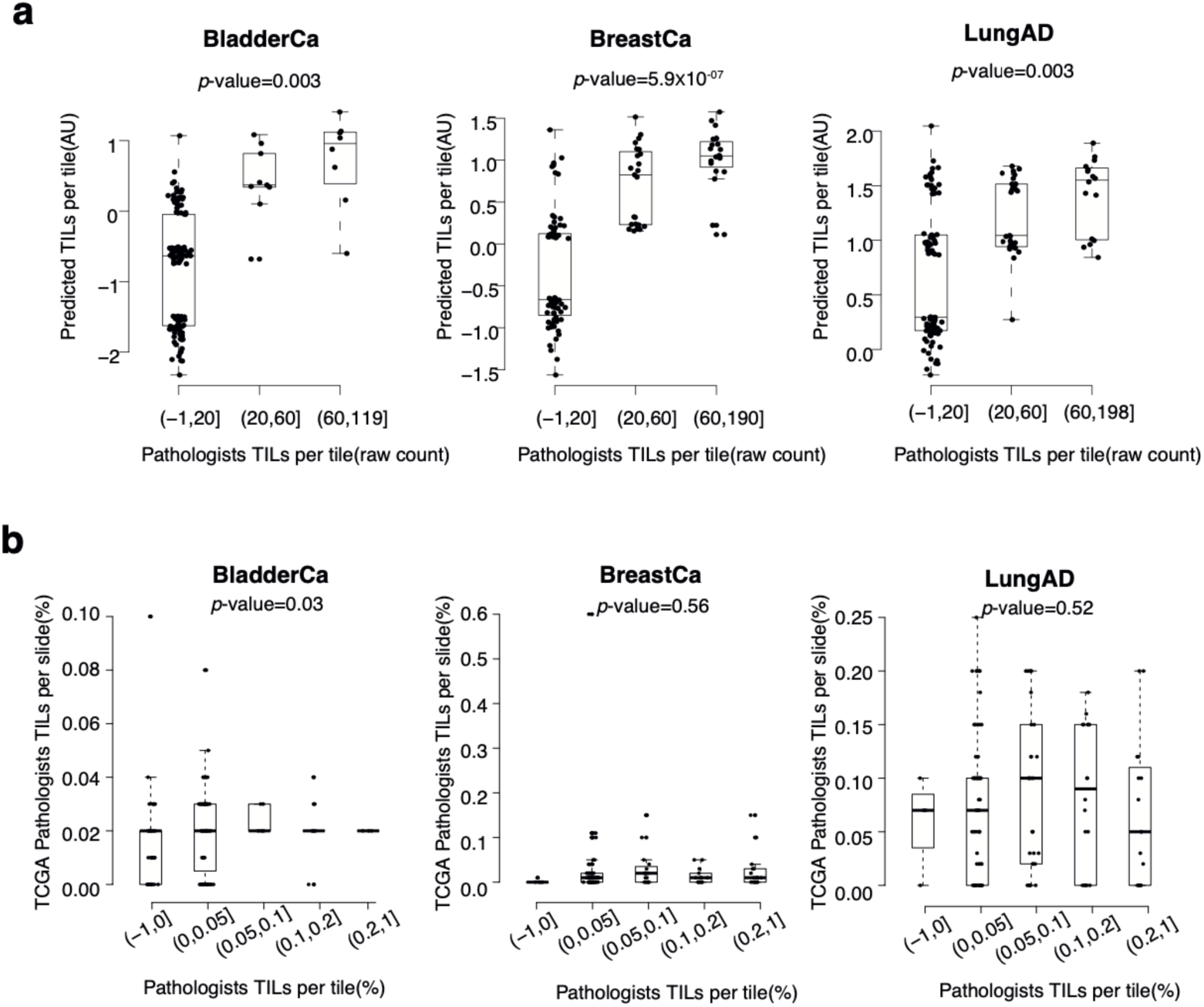
TIL score predicted by PC-CHiP yield higher consistency with histopathologists than slide level TCGA evaluation. **a**, Systematic blinded assessment of TIL raw counts by two expert pathologists for three different cancer types. Each box plot shows the predicted TIL scores from PC-CHiP for tiles with different TIL raw counts, as independently evaluated by pathologists. **b**, slide level evaluation of TILs yield lower concordance with systematic blinded assessment of TIL. Each box plot shows the slide level TILs evaluation from TCGA for tiles with different TIL raw counts.

**Extended Data Figure 7.**
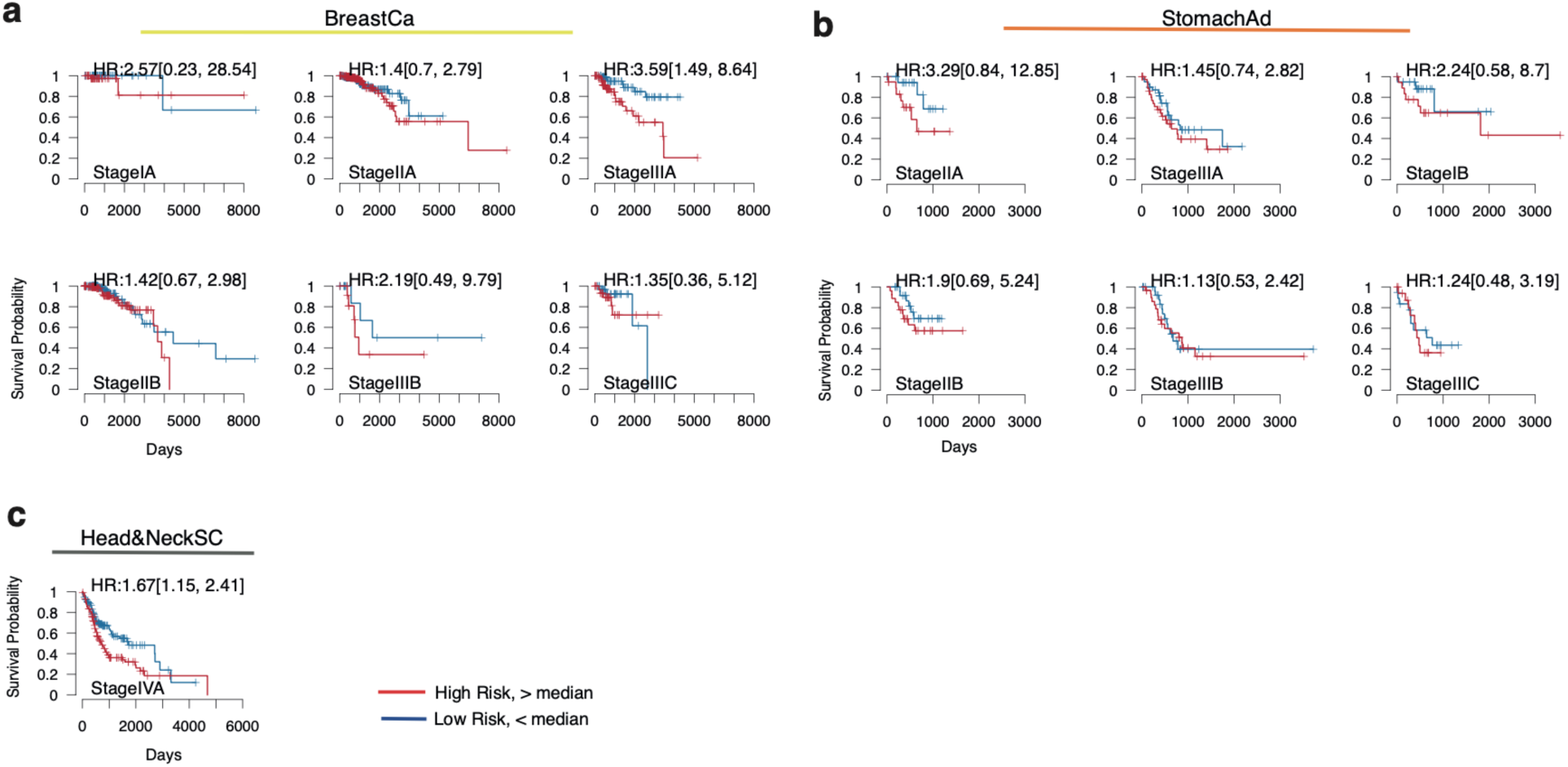
Patient risk stratification using histopathological features. Kaplan-Meier curves for high and low risk groups in different tumor types and stages. **a**, breast invasive carcinoma. **b**, stomach adenocarcinoma. **c**, head and neck squamous cell carcinoma. Only tumor stages with at least 20 patients are shown. Hazard ratios and the corresponding 95% confidence interval were computed using a Cox proportional hazards model.

**Extended Data Figure 8.**
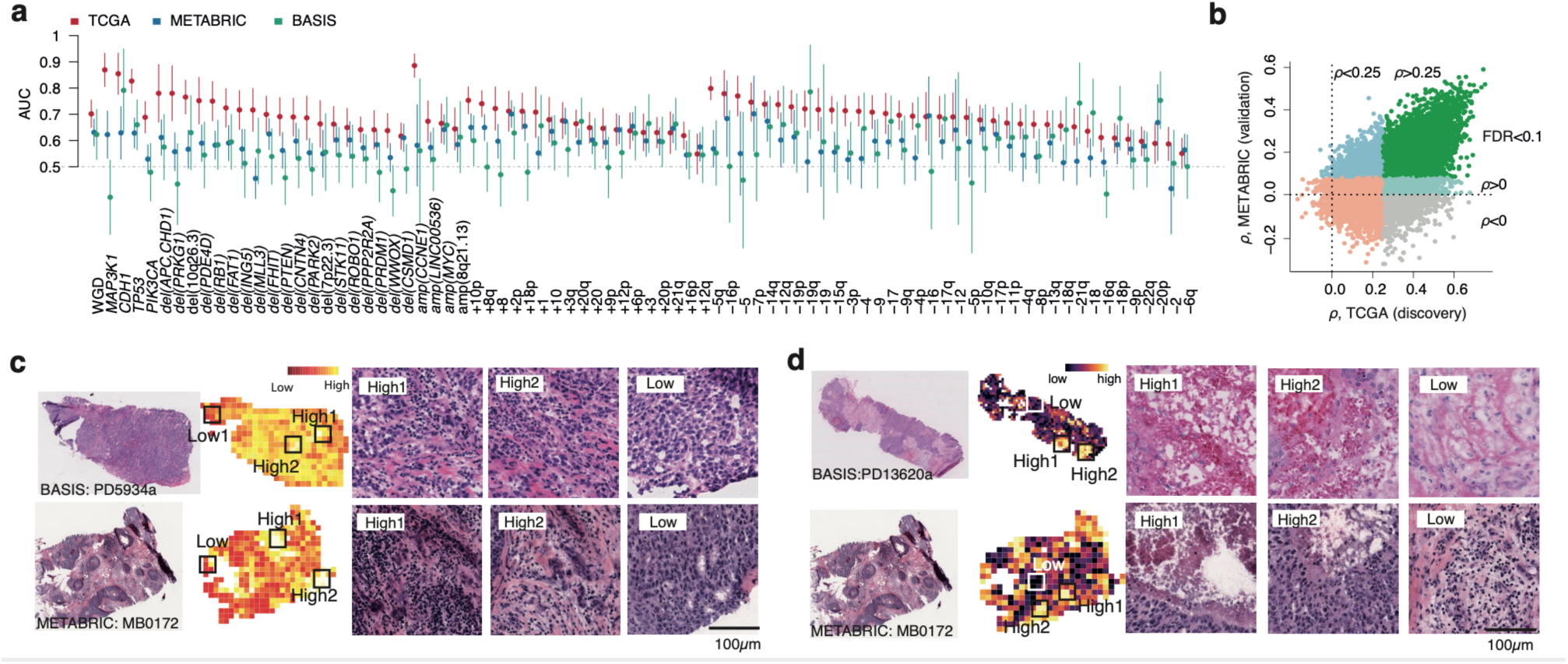
Overall performance of PC-CHiP in validation datasets. **a**, The validation accuracy in METABRIC (blue) and BASIS (green) datasets compared to TCGA dataset (red) for each significant association discovered in TCGA indicated at the bottom. **b**, The distribution of correlation between predicted and true transcript level in METABRIC (x-axis) compared to those in TCGA (y-axis). Each dot represents a gene; blue dots are the genes that can be validated in METABRIC (correlation>0 and FDR<0.1). **c**, Example slides with spatial prediction of TILs. From left to right are the original H&E stained slide, spatial prediction of TILs based on histopathological features, tile predicted as high infiltration and low infiltration. **d**, Example slides with spatial risk prediction. From left to right are the H&E stained slide, the corresponding spatial risk prediction and a selection of enlarged tiles with estimated high or low risk.

**Extended Data Figure 9.**
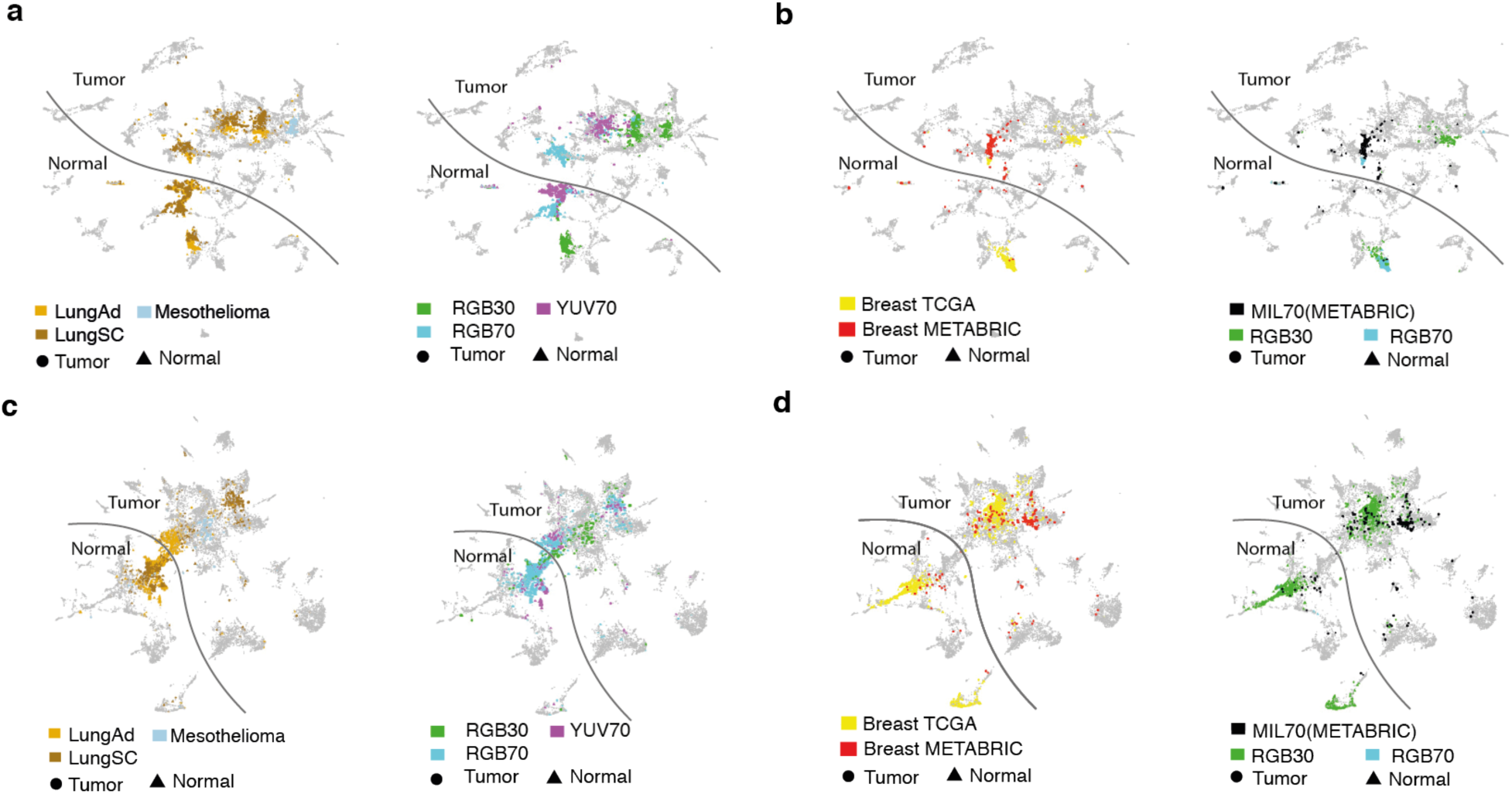
Impact of jpeg quality on histopathological feature representations before and after retraining of Inception-V4. UMAP representation of the histopathological features from the original Inception model (**a, b**) and the modified, retrained architecture (**c, d**). **a**, lung adenocarcinoma, squamous cell carcinoma and normal lung tissue highlighted. **b**, breast tumor and normal from TCGA and breast tumor from METABRIC highlighted. **c**, as in **a**, but for the modified architecture. **d**, as **c** based on the modified architecture. In each figure, the plot on the right side is colored by tissue type and the plot on the left side is colored by jpeg quality.

**Extended Data Figure 10.**
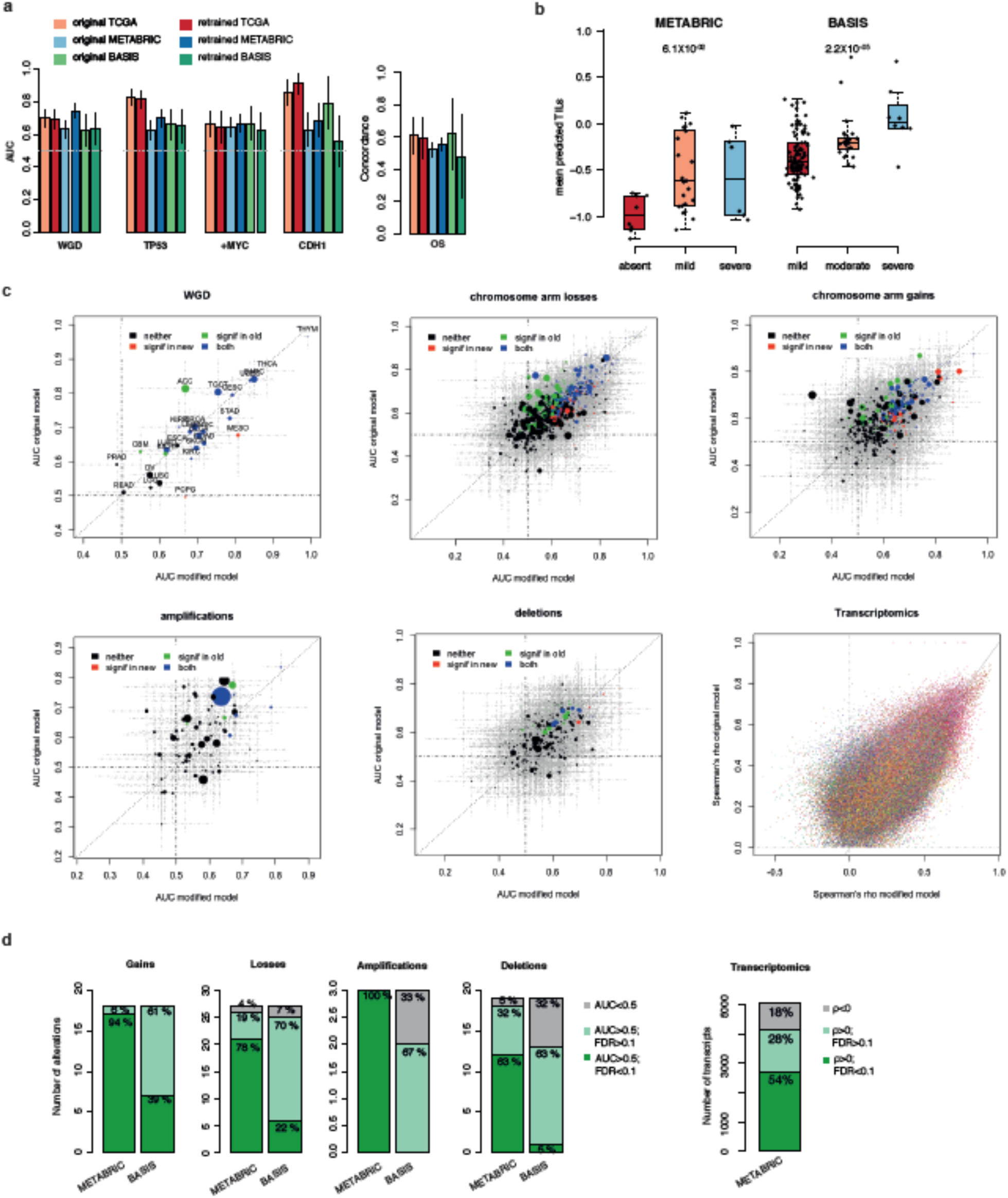
Molecular association of the modified Inception architecture. **a**, AUC for selected genetic alterations and survival for the original and modified Inception architecture. Error bars denote 95% confidence intervals. **b**, Whole-slide average histopathology predictions for TILs from the modified network (*x*-axis) relative to expert pathologist categories (*y*-axis). Boxplots depict the quartiles and median, whiskers extend to 1.5× the inter quartile range. **c**, Scatterplots of genomic and transcriptomic association strengths based on the original (*x*-axis) and modified (*y*-axis) Inception model. Predictions from the original model are five-fold cross-validated, while those of the modified architecture are evaluated on a single 70% training/ 30% testing split. **d**, Distribution of validated (deep green), indeterminate (light green) and invalid (gray) associations in METABRIC and BASIS across different alteration types. Distribution of validated (deep green), indeterminate (light green) and invalid (gray) transcriptomic associations in METABRIC.

## Supplementary Tables

**Supplementary Table 1. Tissue classification performance.**

This table contains 3 sheets. **CancerTypeAbbreviation:** The abbreviation of the TCGA studies in this paper. **TissueClassification:** The tissue classification accuracy in the held-back validation set for each cancer type (by row). First 3 columns correspond to the tissue classification accuracy while considering all 42 tissue types. Last 3 columns correspond to tissue classification accuracy while considering only the normal and tumor tissue of that cancer type. **HistoSubtypeClassification:** The histological subtype classification accuracy for lung and kidney cancers. Accuracies are reported in AUC, the 95% confidence interval was computed using bootstrap.

**Supplementary Table 2. Genetic and transcriptomic associations performance.**

This table contains 11 sheets. In **WholeGenomeDuplication, WGDfromOtherCancers, PointDriverMutation, FocalAmp, FocalDel and ChromArmGain, ChromArmLoss**, the predictive accuracy (AUC) is reported for each cancer type (by row) of the genetic alterations that correspond to the sheet name. The columns from left to right are: lower bound of AUC at 95% confidence level, AUC estimated from 5-CV, higher bound of AUC at 95% confidence level, p value estimated from 5-CV, total number of samples, total number of samples with alteration, alteration name, cancer type and FDR. **Transcription:** The list of significant transcription/cancer associations (Spearman’s *ρ*>0.25, FDR<0.1). From left to right columns are: lower bound of correlation, correlation from 5-CV, higher bound of correlation, p value estimated from 5-CV, FDR, gene name and cancer type. **EnrichedPathway:** the list of significantly enriched (FDR<0.1) pathways among the significantly associated transcripts. From left to right columns are: enriched score, normalised enriched score, associated *p*-value, odds ratio, cancer type, pathway name, pathway function and the adjusted *p*-value using FDR (all statistics computed using GSEA R package). **ProliferationScore and TILs**, the Spearman’s *ρ* of the predicted score from 5-CV for each cancer type. From left to right are cancer type, lower bound of correlation, correlation from 5-CV, higher bound of correlation, p value estimated from 5-CV, FDR.

**Supplementary Table 3. Survival analysis performance.**

**Concordance CV:** The average predicted concordance of overall survival over 5-fold cross-validation in the held-back validation dataset for models using different covariates (each column). From left to right columns are for models using: histopathology (subtypes and grade), PC-CHiP, clinical data (histopathology, stage, gender, age), clinical data and PC-CHiP (retrained with boosting), clinical data and PC-CHiP (pretrained linear predictor), all (clinical + gene expression), all + PC-CHiP (retrained with boosting), all + PC-CHiP (pretrained linear predictor). **Wald Test / Likelihood ratio Test PC-CHiP:** Concordance for models using clinical data (not cross-validated), Wald’s *p*-values for including a pretrained cross-validated linear predictor of PC-CHiP, log-likelihood for the models and likelihood ratio test p-values. **Metrics Survival Models:** The number of covariates available (by column) for each cancer type (by row).

**Supplementary Table 4. Performance of features from amended CNN compared to original CNN. TissueClassif.** The predicted tissue types for METABRIC and BASIS from original CNN (old) and amended CNN (new). Tissue types indicated in row. **ValidationAUC.** Predictive accuracy for alterations that were significant in TCGA (by row) for models built with features from original and amended CNN.

## Supplementary Data

**Supplementary Data 1.**

High resolution images tiles at 20x magnification, 512px x 512px (0.5µm per pixel), shown in Figures 1–4.

